# Subunit redundancy within the NuRD complex ensures fidelity of ES cell lineage commitment

**DOI:** 10.1101/362988

**Authors:** Thomas Burgold, Michael Barber, Susan Kloet, Julie Cramard, Sarah Gharbi, Robin Floyd, Masaki Kinoshita, Meryem Ralser, Michiel Vermeulen, Nicola Reynolds, Sabine Dietmann, Brian Hendrich

## Abstract

Multiprotein chromatin remodelling complexes show remarkable conservation of function amongst metazoans, even though components present in invertebrates are often present as multiple paralogous proteins in vertebrate complexes. In some cases these paralogues specify distinct biochemical and/or functional activities in vertebrate cells. Here we set out to define the biochemical and functional diversity encoded by one such group of proteins within the mammalian Nucleosome Remodelling and Deacetylation (NuRD) complex: Mta1, Mta2 and Mta3. We find that, in contrast to what has been described in somatic cells, MTA proteins are not mutually exclusive within ES cell NuRD and, despite subtle differences in chromatin binding and biochemical interactions, serve largely redundant functions. Nevertheless, ES cells lacking all three MTA proteins represent a complete NuRD null and are viable, allowing us to identify a previously undetected function for NuRD in maintaining differentiation trajectory during early stages of lineage commitment.

## Introduction

Mammalian cells contain a number of proteins capable of using ATP hydrolysis to shift nucleosomes relative to the DNA sequence, thereby facilitating chromatin remodelling. In mammals, these ATP-dependent chromatin remodelling proteins usually exist within multiprotein complexes and play essential roles in the control of gene expression, DNA replication and repair (Hargreaves and Crabtree 2011; Narlikar et al. 2013; Hota and Bruneau 2016).

NuRD (Nucleosome remodelling and deacetylation) is one such multiprotein complex which is unique in that it contains both chromatin remodelling and protein deacetylase activity. NuRD is highly conserved amongst metazoans and has been shown to play important roles in cell fate decisions in a wide array of systems (Denslow and Wade 2007; Signolet and Hendrich 2015). For example, in embryonic stem (ES) cells NuRD controls nucleosome positioning at regulatory sequences to finely tune gene expression (Reynolds et al. 2012; Bornelöv et al. 2018) and in somatic lineages NuRD activity has been shown to prevent inappropriate expression of lineage-specific genes to ensure fidelity of somatic lineage decisions (Denner and Rauchman 2013; Knock et al. 2015; Gomez-Del Arco et al. 2016; Loughran et al. 2017). It was recently demonstrated that this is achieved in both ES cells and B-cell progenitors by restricting access of transcription factors to regulatory sequences (Liang et al. 2017; Loughran et al. 2017; Bornelöv et al. 2018). Additionally, aberrations in expression levels of NuRD component proteins are increasingly being linked to cancer progression (Lai and Wade 2011; Mohd-Sarip et al. 2017).

NuRD is comprised of two enzymatically and biochemically distinct subcomplexes: a chromatin remodelling and a deacetylase subcomplex. The chromatin remodelling subcomplex contains a nucleosome remodelling ATPase protein (Chd3/4/5) along with one of the zinc finger proteins Gatad2a/b and the Doc1/Cdk2ap1 protein, while the deacetylase subcomplex contains class I histone deacetylase proteins Hdac1/2, the histone chaperones Rbbp4/7, the Metastasis Tumour Antigen family of proteins, Mta1, Mta2 and Mta3 and, in pluripotent cells, the zinc finger proteins Sall1/4 (Lauberth and Rauchman 2006; Allen et al. 2013; Kloet et al. 2015; Bode et al. 2016; Low et al. 2016; Miller et al. 2016; Spruijt et al. 2016; Zhang et al. 2016). These two subcomplexes are bridged by Mbd2/3, creating intact NuRD. While HDAC and RBBP proteins are also associated with other chromatin modifying complexes, the CHD, MBD, GATAD2 and MTA proteins are obligate NuRD components. Functional and genetic data indicate that the CHD4-containing remodelling subunit may be capable of functioning independently of intact NuRD (O’Shaughnessy and Hendrich 2013; O’Shaughnessy-Kirwan et al. 2015), though it is not clear whether the deacetylase subcomplex has any function outside of intact NuRD.

Changes in subunit composition in large multiprotein, chromatin modifying complexes such as PRC1 and BAF has been shown to correlate with distinct changes in function to sites of action in the chromatin in a cell-type specific manner (Ho and Crabtree 2010; Morey et al. 2012). The NuRD complex might therefore be expected to show similar diversity in both composition and function and in fact, diversification of NuRD function has been described through differential incorporation of different isoforms of NuRD component proteins (Bowen et al. 2004). For example, Mbd2 and Mbd3 are mutually exclusive within NuRD (Le Guezennec et al. 2006). While Mbd2 is not required for mammalian development, Mbd3 is essential for early postimplantation mouse development (Hendrich et al. 2001). Mbd2/NuRD is a methyl-CpG binding co-repressor complex which is dispensable for early development but Mbd3/NuRD, a transcriptional modulator found at sites of active transcription, has been shown to play important roles in regulation of cell fate decisions in multiple developmental systems (Feng and Zhang 2001; Reynolds et al. 2012; Gunther et al. 2013; Reynolds et al. 2013; Shimbo et al. 2013; Menafra et al. 2014). In brain development NuRD complexes containing either CHD3, CHD4 or CHD5 play distinct roles during cortical development (Nitarska et al. 2016).

Further functional and biochemical diversification occurs through alternate use of the three MTA proteins within NuRD. MTA proteins function as a scaffold around which the deacetylase subcomplex is formed, comprising a 2:2:4 stoichiometry of MTAs:HDACs:RBBPs (Millard et al. 2013; Smits et al. 2013; Millard et al. 2016; Zhang et al. 2016). The three MTA proteins are highly conserved, differing from each other predominantly at their C-termini. The MTA1 protein was originally identified because of its elevated expression in metastatic cell lines (Toh et al. 1994), and subsequently all three MTA proteins have been shown to be up-regulated in a range of different cancer types (Covington and Fuqua 2014; Sen et al. 2014; Ma et al. 2016). The MTA1 and 3 proteins were shown to form distinct NuRD complexes in breast cancer cells and in B-cells and were recruited by different transcription factors to regulate gene expression (Mazumdar et al. 2001; Fujita et al. 2003; Fujita et al. 2004; Si et al. 2015). These studies did not detect biochemical interactions between MTA3 and the other MTA proteins, leading to the conclusion that MTA proteins are mutually exclusive within NuRD. In contrast, Mta1 was shown to interact with Mta2 in MEL cells, possibly indicating that mutual exclusivity may be cell type-specific (Hong et al. 2005). While all three *Mta* genes are expressed in ES cells, detailed biochemical analysis of interactions of MTA proteins with one another or with the various NuRD components in ES cells has not previously been described.

Functional evidence does not support a strict lack of redundancy amongst MTA proteins during mammalian development. While zygotic deletion of *Chd4* or *Mbd3* results in pre- or peri-implantation developmental failure respectively (Kaji et al. 2007; O’Shaughnessy-Kirwan et al. 2015), mice deficient in any one of the three MTA proteins show minimal phenotypes. Mice lacking either *Mta1* or *Mta3* are viable and fertile (Manavathi et al. 2007)(Mouse Genome Informatics), while mice lacking *Mta2* show incompletely penetrant embryonic lethality and immune system defects (Lu et al. 2008). In the current study we took a systematic approach to dissecting MTA protein biochemical and functional diversity. We find that, in contrast to what has been described in somatic cells, MTA proteins are not mutually exclusive within NuRD in ES cells and serve largely redundant functions. Furthermore, ES cells lacking all three MTA proteins are viable and represent a complete NuRD null, allowing us to identify a previously undetected function for NuRD in early stages of lineage commitment.

## Results

### Mta proteins are not mutually exclusive within the NuRD complex in ES cells

The absence of a detected interaction between the MTA2 and MTA3 proteins in human cells (Fujita et al. 2003; Si et al. 2015), and the observation that different MTA proteins can show different protein-protein interactions in B-cells (Fujita et al. 2004) has led to the conclusion that the MTA proteins are mutually exclusive within NuRD, and could hence confer functional diversity to the NuRD complex (Lai and Wade 2011). To investigate the biochemical specificity of the MTA proteins in an unbiased manner, we used gene targeting of endogenous loci to produce three different mouse ES cell lines in which an epitope tag was fused to the C-terminus of each MTA protein (Fig. 1A). Although MTA genes show different expression patterns in preimplantation mouse development, all three are expressed in peri-implantation and early postimplantation epiblast, the tissue most similar to the naïve ES cell state (Fig. S1). We therefore considered ES cells to be a good system in which to investigate the function of MTA proteins.

**Figure 1.**
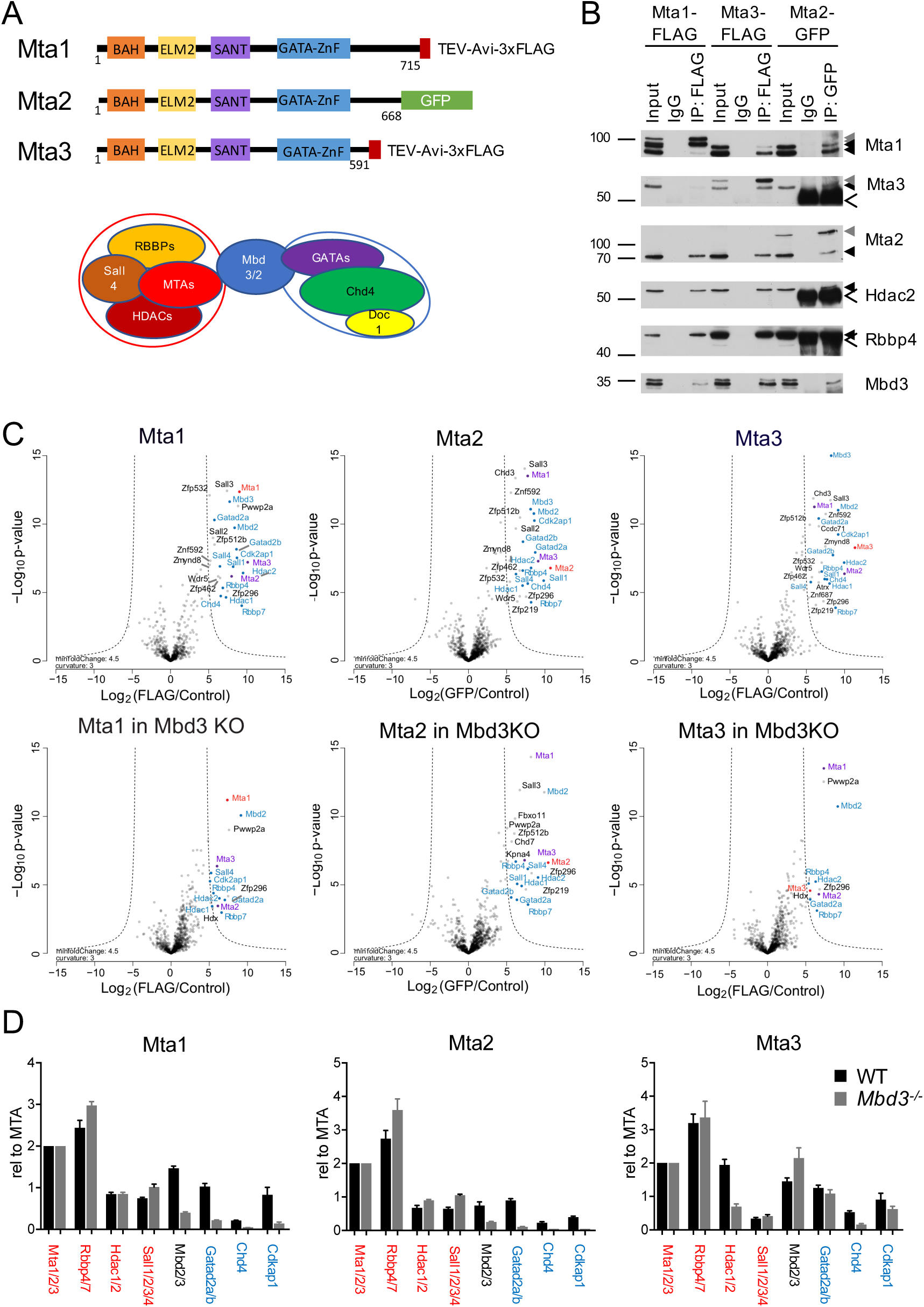
Biochemical characterisation of epitope-tagged MTA proteins. A. Schematic of MTA proteins with different protein domains indicated as coloured boxes, and the C-terminal epitope tags indicated. B. Tagged MTA proteins in heterozygously targeted cell lines were immunoprecipitated and subsequent western blots probed with antibodies indicated at right. Solid black triangles indicate the locations of untagged proteins, grey triangles show the position of epitope-tagged proteins, and open arrowhead show the locations of IgG bands in the final two lanes. Molecular weight in KDa are shown at left. (Full western blot images are available in Mendeley Data.) C. Proteins co-purifying with Mta1, Mta2 or Mta3 in IP-mass spectrometry experiments in wild type cells (top) or Mbd3-null cells (bottom). Proteins showing significant enrichment with the bait protein are located outside the dotted lines. For all panels the protein being immunoprecipitated is indicated in red, the other MTA proteins in purple, and other NuRD components in blue. Each IP/Mass spec experiment was carried out in biological triplicate. D. Relative enrichment of indicated proteins in MTA pulldowns from wild type (WT) or *Mbd3-null* (Mbd3KO) ES cells, normalised to 2x MTA proteins. NuRD components comprising the remodelling subunit are labelled in blue, those comprising the deacetylase subunit in red.

Each tagged protein was expressed at levels comparable to those of wild type proteins and was found to interact with other NuRD component proteins by immunoprecipitation (Fig. 1B). Each MTA protein was also able to immunoprecipitate both of the other MTA proteins in addition to unmodified forms of itself (Fig. 1B). Each NuRD complex contains two copies of an MTA protein (Millard et al. 2013; Smits et al. 2013; Zhang et al. 2016), so the identification of an interaction between MTA proteins means that individual NuRD complexes in ES cells could contain either homodimers or heterodimers consisting of any combination of the three MTA proteins.

To investigate the potential biochemical diversification conferred by different MTA proteins in pluripotent cells we used our tagged cell lines to identify proteins interacting with each of the MTA proteins by label-free quantitative mass spectrometry. Each protein robustly co-purified with all known NuRD component proteins, including each of the MTA proteins, confirming that MTA proteins are not mutually exclusive within NuRD in ES cells (Fig. 1C). In addition to NuRD components, each of the MTA proteins co-purified with Wdr5 as well as a number of zinc finger proteins, most of which had previously been identified as NuRD-interacting proteins (Bode et al. 2016; Spruijt et al. 2016; Ee et al. 2017; Matsuura et al. 2017).

As NuRD is assembled from a deacetylase subcomplex and a remodeller subcomplex joined through the Mbd3 protein (Fig. 1A), loss of Mbd3 is expected to result in dissociation of these two subcomplexes. Endogenous tagging for each Mta protein was performed in an ES cell line harbouring a floxed *Mbd3* allele, and IP/Mass spectrometry was repeated in ES cells after *Mbd3* deletion in order to enrich for interactions specific for the deacetylase subcomplex. The majority of interactions with non-NuRD components was lost after *Mbd3* deletion, indicating that most of these proteins do not directly associate with either the MTA proteins or with the deacetylase subcomplex (Fig. 1C, Table S1). Exceptions to this were Zfp296, which was identified as interacting with all three MTA proteins in an *Mbd3-*independent manner, and Pwwp2a, which co-purified with Mta1 in both wild type and Mbd3-null cells. Notably, Zfp219 and Zfp512b both showed Mbd3-independent interactions specifically with Mta2 (Fig. 1C, Table S1).

By quantitating the abundance of peptides sequenced in each experiment we found that the interactomes for both Mta1 and Mta2 showed a depletion of peptides associated with the remodeller subcomplex (i.e. Chd4, Gatad2a/b and Cdk2ap1) in *Mbd3*-null cells, whereas proteins associated with the histone deacetylase subcomplex (Mta proteins, Hdac1/2, Sall proteins and Rbbp4/7) remained present at similar levels (Fig 1D). In contrast the Mta3 interactome showed an increased interaction with Mbd2 and no relative loss of either sub-complex in the absence of Mbd3. Thus both Mta1 and Mta2 can form part of a stable deacetylase subcomplex in the absence of intact NuRD, but Mta3 is preferentially found in intact NuRD complexes. All three MTA proteins associated with Mbd2 in the absence of Mbd3, indicating that they can all can contribute to Mbd2/NuRD. Together these data show that the MTA proteins are found exclusively within the NuRD complex in ES cells.

### MTA proteins show subtle differences in genome-wide binding patterns

To test whether MTA proteins confer differential chromatin binding to NuRD complexes we subjected our ES cell lines expressing epitope-tagged MTA proteins to chromatin immunoprecipitation followed by high throughput sequencing (ChIP-Seq). The binding profiles of the three proteins were largely, but not completely overlapping (Fig. 2A). Mta3 peaks were almost entirely associated with Mta1 and/or Mta2-binding, while 31% of Mta1 peaks and 22% of Mta2 peaks were not associated with any other MTA protein (Fig. 2A). The vast majority of MTA peaks overlapped with Chd4 peaks, indicating that they represent NuRD-bound regions (Figs. 2B, C). Mta3 peaks were almost exclusively associated with Chd4 binding, consistent with Mta3 being preferentially in intact NuRD (Fig. S2A). In contrast, 15% of Mta1 peaks and 10% of Mta2 peaks did not overlap with the Chd4 dataset.

**Figure. 2.**
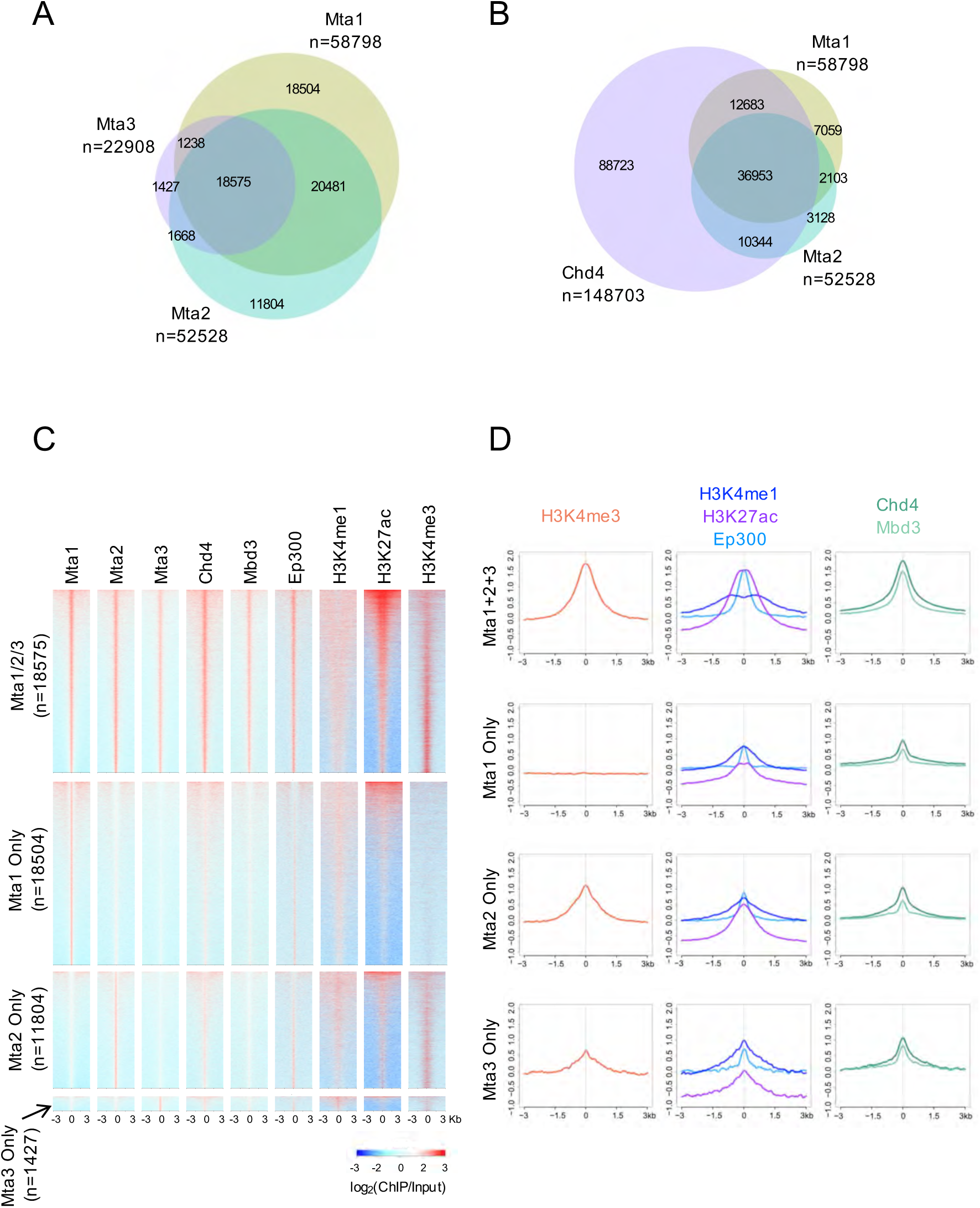
Mta proteins show similar chromatin binding patterns. A. Comparison of peaks identified by ChIP-seq for each MTA protein in wild type ES cells. Total peak numbers are indicated below each protein name. Each ChIP-seq dataset was made from biological triplicates. B. Comparison of Mtal and Mta2 peaks with Chd4 peaks. C. ChIP-seq enrichment for indicated proteins or histone modifications is plotted across different subsets of Mta-bound sites. Mta1/2/3 refers to peaks identified in all three ChIP-seq datasets, whereas “Mta1 Only”, “Mta2 Only” or “Mta3 Only” refer to peaks only called for that protein. D. Average enrichment of density plots in C are plotted for different subsets of Mta ChlP-seq peaks.

Sites found associated with all three MTA proteins were highly enriched for Chd4 and Mbd3 binding, consistent with these being core NuRD binding regions (Fig. 2C, top panels). These sites were also enriched for marks of active promoters (H3K4Me3, H3K27Ac) and active enhancers (H3K4Me1, H3K27Ac, P300; Fig. 2C, D), both of which are hallmarks of NuRD-associated regions (Miller et al. 2016; Bornelöv et al. 2018). The same was true for sites bound by any two of the three MTA proteins (Fig. S2B). The majority of sites occupied by one MTA protein, but not the other two, were also occupied by Chd4 but to a lesser extent than is seen at core NuRD sites (Fig. 2C, D). Whereas Mta2-only and Mta3-only sites showed enrichment for both enhancer and promoter marks, Mta1-only sites showed no enrichment for H3K4Me3, indicating that Mta1 is not found at promoters either without one of the other Mta proteins or within the context of complete Mbd3/NuRD (Fig. 2C, D, S2C). In contrast, sites bound by Mta2 and Mta3, but not Mta1 showed particular enrichment for H3K4Me3 (Fig. S2B). None of the MTA-bound sequences showed any enrichment for methylated DNA (Fig. S2D). In all cases MTA-only sites lacking Chd4, which could represent sites of binding by the histone deacetylase subcomplex only, showed characteristics of inactive enhancers, in that they were moderately enriched for H3K4Me1 and P300, but not for H3K4Me3, H3K27Ac or H3K36Me3 (Fig S2C). Genes associated with binding by only one or two MTA proteins, with or without Chd4 are associated with a similar distribution of GO terms as core NuRD sites (Fig S2E) which is not consistent with the idea that the MTA proteins are directing NuRD activity to specific gene subsets.

### Mta1/Mta2/Mta3 triple knockout is a total NuRD null

To identify specific functions for the different MTA proteins, we obtained gene trap alleles for each of the three MTA genes from the European Conditional Mouse Mutagenesis Programme (Skarnes et al. 2011) as ES cell lines (*Mta1* and *Mta2*) or as embryos (*Mta3*) (Fig. S3A-C). ES cell lines were used for morula aggregation to create chimaeric mice which were subsequently outcrossed to wild type (C57Bl/6) mice to establish a mouse line. Conditional deletion alleles were generated for each line, which was subsequently bred with females expressing Sox2-Cre (Hayashi et al. 2002) resulting in deletion of floxed exons and the absence of any detectable protein production from each allele (Fig. S3D; see Methods for Burgold et al. Fig 3 details). As has been reported previously, *Mta1^-/-^* and *Mta3^-/-^* mice were viable, and Mta2^-/-^ mice showed incompletely penetrant embryonic lethality (Manavathi et al. 2007; Lu et al. 2008).

**Figure 3.**
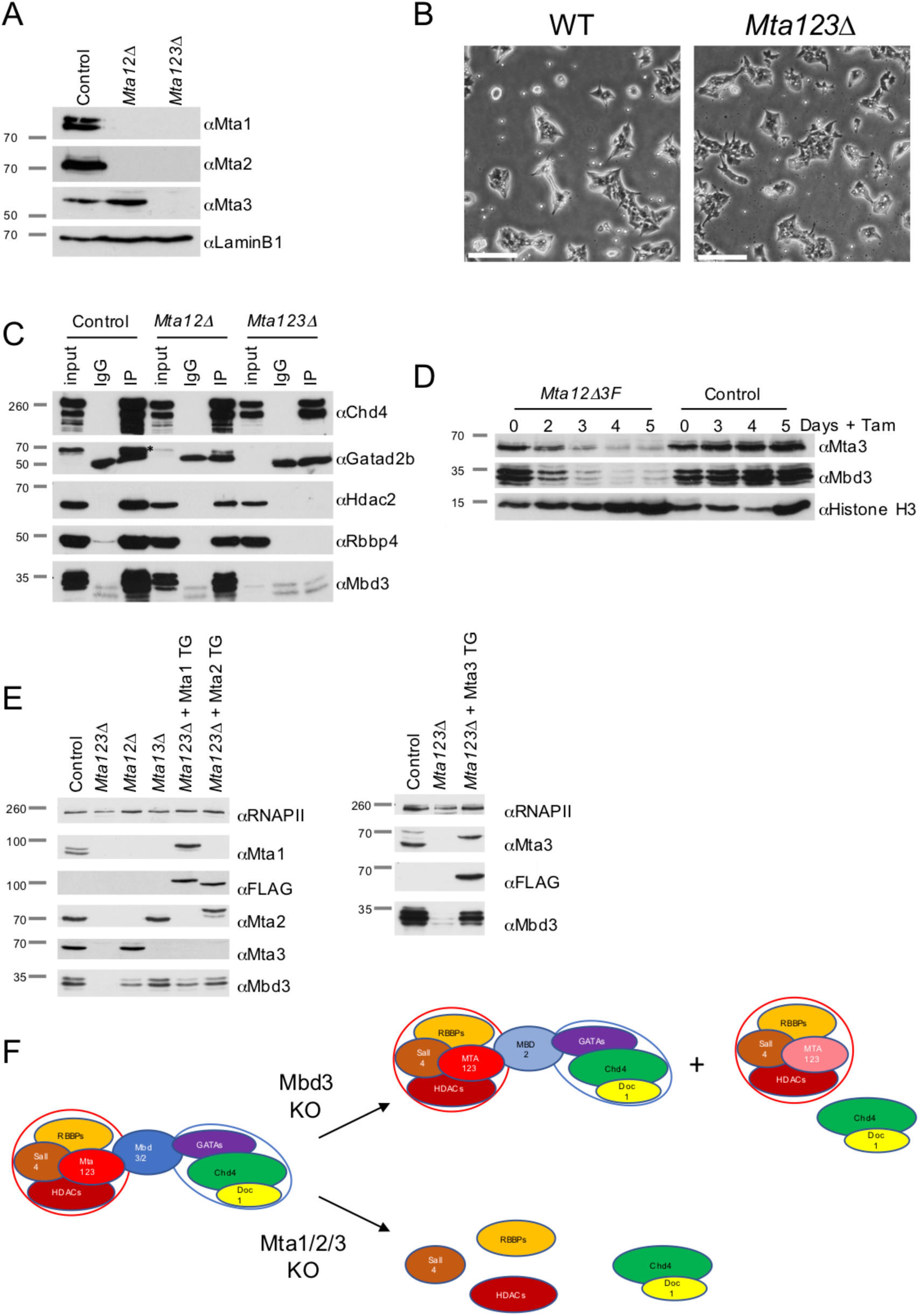
Mta1/Mta2/Mta3 triple null ES cells represent a complete NuRD KO. A. Western blots of wild type (Control), double or triple-null ES cell nuclear extract was probed with antibodies indicated at right. LaminB1 acts as a loading control. Approximate sizes are indicated in KDa. B. Phase-contrast images of wild type or *Mta123Δ* ES cells in self-renewing conditions. Scale bars represent 100μm. C. Western blots of anti-Chd4 immunoprecipitation of nuclear extract from indicated cell lines (top) probed with antibodies indicated at right. Approximate sizes are indicated in KDa. D. Western blot of a time course of *Mta3* deletion in *Mta1ΔMta2ΔMta3^Flox/Flox^: Cre-ER* (Mta12Δ3F) or Control (Mta12Δ3F without Cre-ER) ES cells probed for Mta3, Mbd3 or Histone H3 as a control. The time course is indicated at the top as Days + tamoxifen. E. Western blot showing rescue of Mta123Δ ES cells by ectopic expression of Mta1, Mta2 or Mta3 from a transgene (TG). Total RNA Polymerase II acts as a loading control (RNAPII). For all western blots molecular weight is indicated at left in KDa. F. Model of NuRD complex structure. Upon loss of Mbd3 most of the NuRD complex falls apart into the Chd4-containing remodeller subcomplex and the MTA-containing deacetylase subcomplex, but some intact Mbd2-NuRD still remains. Upon loss of all three MTA proteins no intact NuRD remains, both Mbd3 and Gatad2a/b become unstable and neither of the intact subcomplexes remain.

*Mta1, Mta2* or *Mta3*-null ES cell lines derived from mice were morphologically indistinguishable from wild type ES cells, as were ES cell lines deficient for combinations of pairs of MTA genes (Fig. S3E). *Mta^-/-^/Mta2^-/-^Mta3^Flox/Flox^* ES cells (*Mta12Δ*) were subsequently created and expanded in culture (see Methods for details; Fig. 3A). After transfection with a Cre expression construct to induce deletion of both *Mta3* alleles we recovered ES cells lacking all three MTA proteins. These *Mta1^-/−/^Mta2^-/-^Mta3^-/-^* ES cells (subsequently referred to as *Mta123Δ*) appeared morphologically normal in standard, 2i + LIF culture (Fig. 3B). MTA proteins are therefore dispensable for ES cell viability.

Structurally, MTA proteins bridge an interaction between the deacetylase subcomplex with Mbd3 and the remodelling subcomplex (Fig. 1A) so we predicted that loss of all three MTA proteins would prevent NuRD formation. Consistent with this prediction we could detect no interactions between Chd4 and components of the deacetylase subcomplex (Hdac2, Rbbp4) in *Mta123Δ* ES cell nuclear extract by immunoprecipitation of endogenous proteins (Fig. 3C). Surprisingly, despite being transcribed at normal levels, both Gatad2b and Mbd3 proteins were barely detectable in *Mta123Δ* cells (Fig. 3C), indicating that the MTAs are important for the stability of both of these proteins. To investigate this further we monitored loss of Mta3 in *Mta1^Δ/Δ^Mta2^Δ/Δ^Mta3^flox/flox^* ES cells after deletion using a tamoxifen-inducible Cre and found that loss of Mbd3 protein stability was coincident with loss of Mta3 protein (Fig. 3D). Furthermore, introduction of either Mta1, Mta2 or Mta3 into *Mta123Δ* ES cells at levels comparable to wild type expression resulted in restoration of Mbd3 protein levels, demonstrating that contact with at least one of the MTA proteins is sufficient for Mbd3 protein stability (Fig. 3E). In contrast to *Mbd3-null* ES cells which display a significant depletion, but not a complete loss of NuRD due to partial compensation by Mbd2 (Kaji et al. 2006), *Mta123Δ* ES cells were completely devoid of any detectable intact NuRD (Fig. 3C, F). The *Mta123Δ* ES cells are thus a total NuRD null ES cell line, which allowed us to examine, for the first time, the consequences of a complete loss of the NuRD complex in a viable mammalian cell system.

### The NuRD complex safeguards cellular identity during differentiation

Twice as many genes showed an increase rather than a decrease in expression levels in *Mta123Δ* ES cells compared to control ES cells by RNA-seq (Fig. 4A). This is consistent with NuRD acting predominantly, but not exclusively, as a transcriptional repressor. The ratio of increased to decreased gene expression in *Mta123Δ* cells was very similar for genes bound by all three Mta proteins, Mta1+2, Mta1 only or Mta2 only (Fig. 4B), indicating that NuRD’s impact on transcription is not detectably altered by the inclusion or absence of individual MTA subunits. Sites bound by MTA proteins in the absence of Chd4, and presumably representing sites bound by the deacetylase subcomplex, were not associated with specific gene misexpression events in the *Mta123Δ* ES cells (Fig. S4A, B).

**Figure 4.**
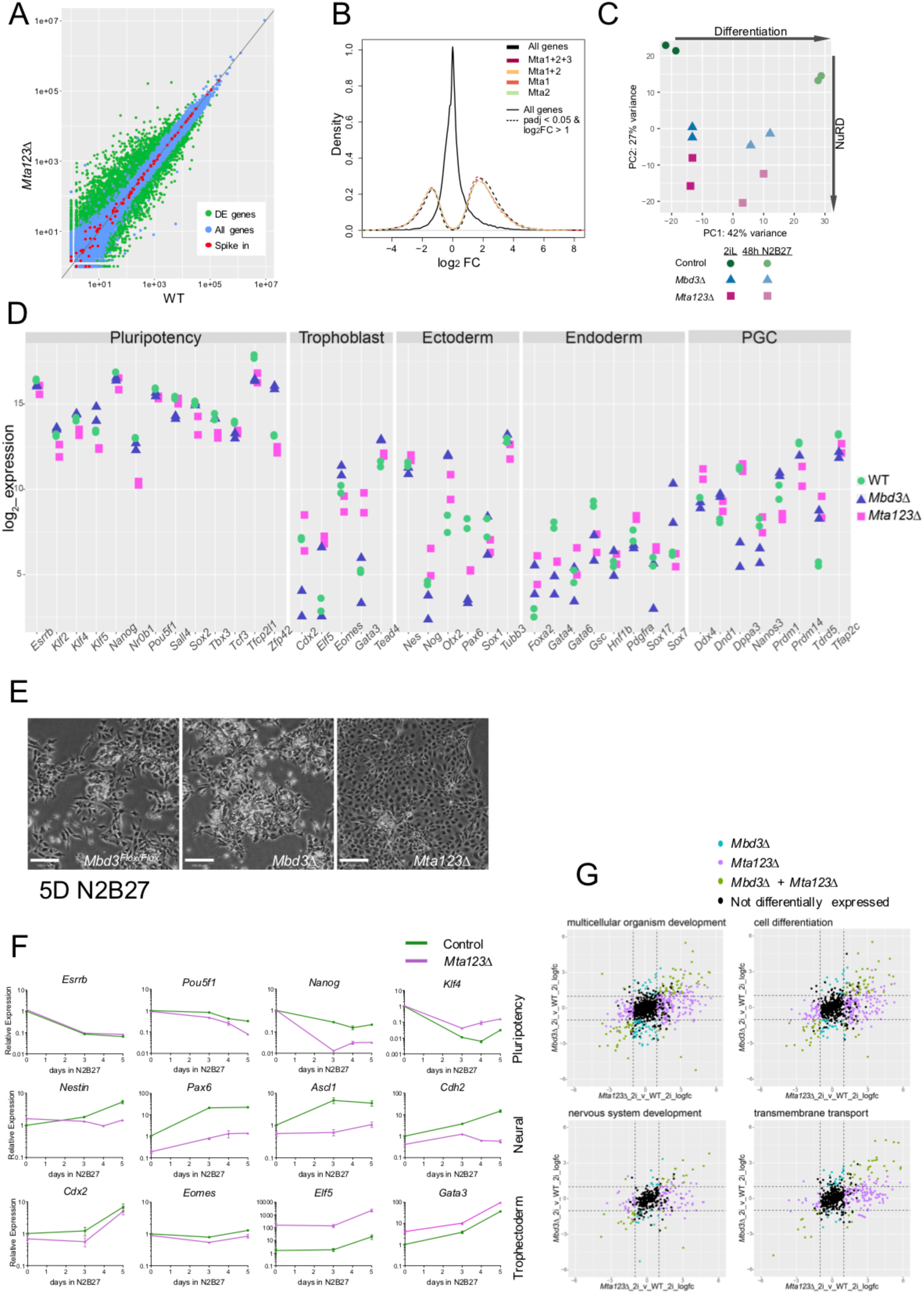
Mta proteins act redundantly to control gene expression. A. Comparison of gene expression in Mta123Δ ES cells and wild type (WT) ES cells. Each circle represents a gene: red indicates spike-in controls, blue indicates genes that are not differentially expressed to a significant degree, and green indicates differentially expressed genes (2404 increased, 1293 decreased) defined with an adjusted p-value < 0.05 and a log2 fold-change > 1. N=3 for each genotype. B. Fold change in gene expression is plotted for different subsets of genes. All genes are plotted in black, while subsets of genes located nearest to ChIP-seq peaks for the indicated proteins +Chd4 and which show significant changes in expression compared to wild type cells are plotted in dashed coloured lines as indicated. The number of genes in each of the Mta categories are: all genes (32271), all differentially expressed (n=3701), Mta1+2+3 (n=1738), Mta1+2 (n=1460), Mta1 (n=1020) and Mta2 (n=924). C. Principal component analysis of RNAseq data from ES cell lines of indicated genotypes in either self-renewing conditions (2i) or after 48 hours in the absence of two inhibitors and LIF (Diff48). Each point represents a biological replicate. D. Expression of indicated genes segregated by functional category for wild type (WT, green circles), *Mbd3Δ* (blue triangles), or *Mta123Δ* (magenta squares). Data are taken from RNA-seq experiments and each point represents the merging of two biological replicates each for 1 (Mbd3Δ) or two (WT, Mta123Δ) independently derived cell lines. E. Phase contrast pictures of ES cells of indicated genotypes after 5 days in differentiation conditions (N2B27). Scale bars represent 100 μm. F. Expression of indicated genes was measured by RT-qPCR in wild type (Control, red), *Mbd3Δ* (purple) or *Mta123Δ* (blue) ES cells over a time course of differentiation. N ≥ 3 biological replicates, error bars indicate SEM. G. Genes associated with indicated GO terms plotted by fold change in expression in *Mta123Δ* ES cells (x-axis) or *Mbd3Δ* ES cells (y-axis) induced to differentiate for 48 hours. Genes are coloured if they are differentially expressed (log2 fold-change > 1 and padj value < 0.05) in either comparison as indicated. The dotted lines show the fold-change cut-off of 2. GO-terms were identified using David v.6.8 (Huang da et al. 2009) using a Benjamini score with a cutoff of 0.05.

Comparing global gene expression profiles using principal component analysis (PCA) showed that *Mta123Δ* and *Mbd3Δ* ES cells in self-renewal conditions were more similar to each other than either was to wild type cells (Fig. 4C). *Mta123Δ* ES cells were most distinct from wild type cells, consistent with them representing a more complete NuRD knockout (Fig. 4C, S4C). Consistent with this, genes driving the NuRD-specific separation (Principal Component 2; PC2) were generally misexpressed to a greater degree in *Mta123Δ* ES cells than in *Mbd3Δ* ES cells (Fig 4D, S4C). As has been shown for *Mbd3Δ* ES cells, both NuRD mutant lines moderately misexpressed pluripotency-associated genes in ES cells (Reynolds et al. 2012)(Fig. 4D). Yet in 2iL conditions pluripotency-associated genes were mis-expressed to a much lesser extent than genes associated with differentiation (Fig. 4D). This demonstrates that, in addition to maintaining pluripotency gene expression levels, NuRD also functions in ES cells to prevent inappropriate activation of lineage-specific gene expression.

Despite showing inappropriate expression of a large number of genes normally associated with differentiated cells, *Mta123Δ* ES cells could be maintained as morphologically undifferentiated ES cells in the very restrictive 2iL conditions (Fig. 3B). We therefore next asked what impact complete loss NuRD activity had upon the differentiation capacity of *Mta123Δ* ES cells. Upon removal of the two inhibitors and LIF from the culture media wild type cells began to adopt the flatter morphology of neurectoderm (Fig. 4E)(Ying et al. 2003). This was accompanied by downregulation of pluripotency-associated genes and the activation of a neural gene expression programme (Fig. 4F, S4C). *Mbd3Δ* ES cells are able to respond to the absence of self-renewal factors but have a very low probability of adopting a differentiated fate when induced to differentiate in N2B27 conditions (Kaji et al. 2006; Reynolds et al. 2012). Consistent with these findings, after 5 days in differentiation conditions *Mbd3Δ* ES cells showed some signs of having responded to differentiation conditions, but still retained pockets of morphologically undifferentiated cells (Fig. 4E). In contrast the completely NuRD-null *Mta123Δ* ES cells appeared to have all exited the self-renewal programme and adopted a flat, monolayer morphology (Fig. 4E). The ability of *Mta123Δ* ES cells to undergo morphologically normal neuroectodermal differentiation was rescued upon re-expression of either Mtal, Mta2 or Mta3 (Fig. S5A).

Since the differentiation process may be considered as a combination of exit from selfrenewal (downregulation of pluripotency-associated genes) and acquisition of lineage specific gene expression programs, we focussed on changes to these two classes of genes by RT-qPCR over a differentiation time course. Pluripotency-associated genes such as Esrrb, *Klf4, Nanog* and *Pou5f1* were downregulated during differentiation of both wild type and *Mta123Δ* ES cells (Fig. 4F). While this downregulation was not absolutely dependent on the presence of functional NuRD, the magnitude and kinetics of the response varied in a gene-dependent manner. Genes associated with acquisition of neural fate were activated in the presence of wild type NuRD, a response which was reduced in the NuRD null line (Fig. 4F). Globally, *Mta123Δ* ES cells misexpressed considerably more genes associated with development and differentiation than did *Mbd3Δ* ES cells after 48 hours of differentiation (Fig. S4E). Exogenous expression of either Mta1, Mta2 or Mta3 in *Mta123Δ* ES cells was able to rescue the ability of cells to activate neural gene expression to different extents during differentiation (Fig. S5B), consistent with their ability to rescue the morphological phenotypes. *Mta123Δ* ES cells induced to differentiate towards a mesoderm fate similarly failed to appropriately activate differentiation markers, and exogenous expression of individual MTA proteins was again able to rescue the defect to varying degrees (Fig. S5C). Together these data show that while NuRD activity is not required for ES cells to exit the naïve state, it is required for activation of lineage-appropriate gene expression as well as to prevent expression of genes associated with other cell types.

To better understand how NuRD-null cells respond when induced to differentiate we compared their gene expression profiles to a transcription landscape made from single cells taken from early mouse embryos (Fig. 5A; S5D) (Mohammed et al. 2017). In the self-renewing state, *Mbd3Δ* and *Mta123Δ* cells clustered near control (parental) lines near embryonic day 3.5 (E3.5) and E4.5 inner cell mass cells, as is expected for naïve mouse ES cells (Boroviak et al. 2014). After 48 hours of differentiation control ES lines clustered with E5.5 epiblast cells, demonstrating that the ES cell differentiation process occurred analogously to development *in vivo*. While *Mbd3Δ* ES cells appear to have exited the self-renewing state and to have taken the same differentiation trajectory as wild type cells (i.e. leftwards along PC1; Fig 5A, S5D), they did not travel as far along this trajectory as wild type cells, instead occupying a space between the E4.5 and E5.5 epiblast states. *Mta123Δ* cells travelled even less far along PC1 than *Mbd3Δ* ES cells, and rather than maintain the appropriate differentiation trajectory they also travelled along PC2, occupying a space between E4.5 epiblast and E4.5 primitive endoderm. This further demonstrates that not only is NuRD important for cells to be able to adopt the appropriate gene expression programme for a given differentiation event, but it is also important for cells to maintain an appropriate differentiation trajectory.

**Figure 5.**
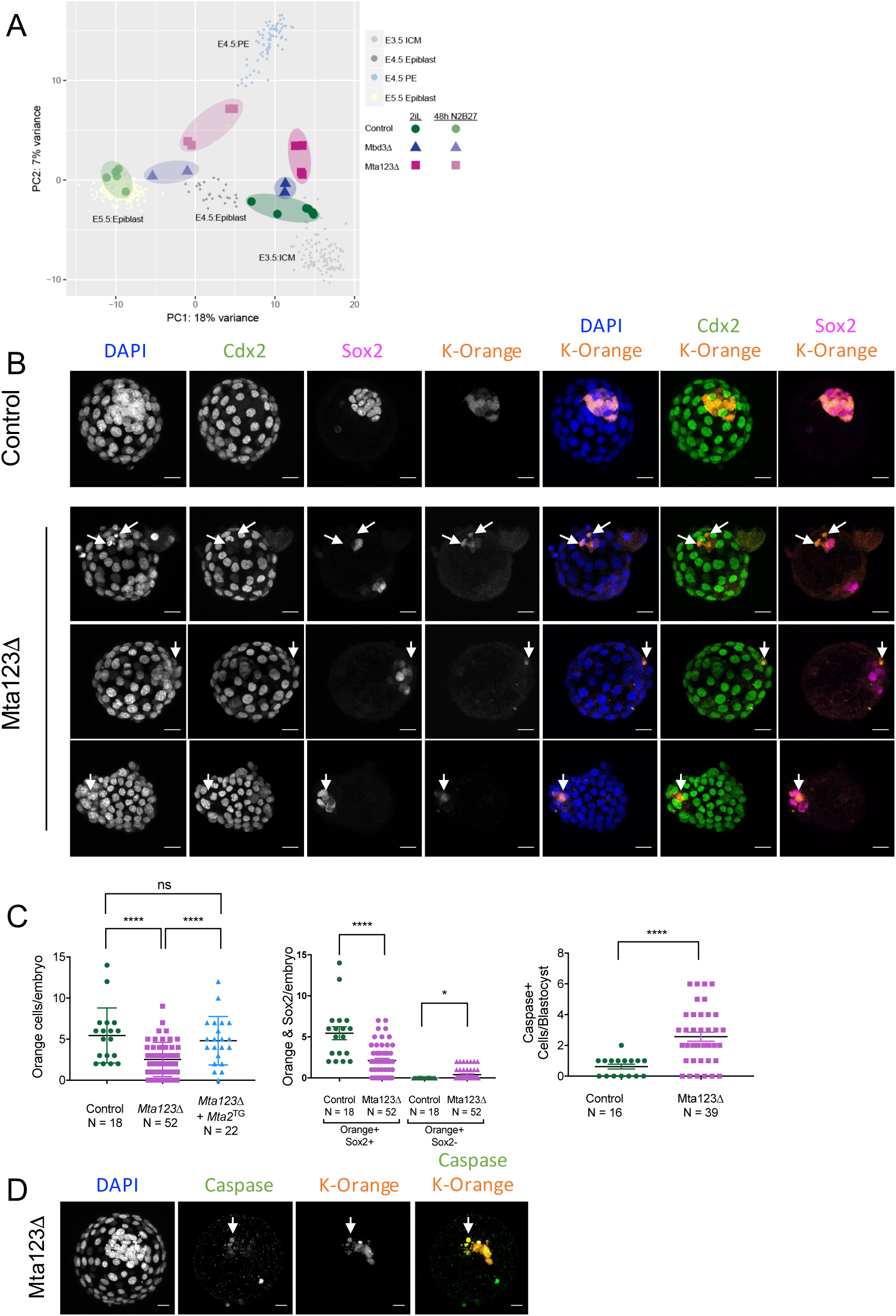
NuRD activity maintains an appropriate ES cell differentiation trajectory. A. Comparison of expression data for wild type, *Mbd3Δ* or *Mta123Δ* ES cells in self-renewing (2i) conditions or after 48 hours in differentiation conditions (Diff48) with mouse embryonic single cell RNA-seq data from (Mohammed et al. 2017). Larger circles represent biological replicates, smaller circles represent individual cells. PC4 vs PC1 is plotted in Fig. S5D. B. Composite images of representative chimaeric embryos made with control (*Mta1^Flox/Δ^Mta2^+/+^Mta3^Flox/Flox^*) or *Mta123Δ* ES cells. ES-derived cells express the Kusabira Orange fluorescent marker. Sox2 indicates epiblast cells and Cdx2 is expressed in trophectoderm cells. Arrows indicate examples of K-Orange expressing cells in the mutant embryos. Scale bars = 20μm. C. (Left) Number of K-Orange expressing cells observed in chimaeric embryos obtained using control ES cells, *Mta123Δ* ES cells, or *Mta123Δ* ES cells in which Mta2 was reintroduced on a constitutively expressed transgene (*Mta123Δ+Mta2^TG^*). P-values calculated using a two-tailed t-test. (Middle) Mean number of K-Orange cells per embryo separated by Sox2 expression. (Right) Number of K-Orange and Caspase-3 positive cells per embryo. P-values calculated using a one-tailed t-test: *P < 0.05, ****P < 0.0001, “ns” = not significant. D. Composite images of representative chimaeric embryos as in Panel B stained with an anti-activated Caspase 3 antibody. Arrows indicate an example of an orange, apoptotic cell. Scale bars = 20μm.

If *Mta123Δ* ES cells were undergoing a specific trans-differentiation event towards trophectoderm during ES cell differentiation then this could become more pronounced if exposed to normal differentiation conditions in a chimaeric embryo. If, in contrast, they are simply unable to differentiate properly, they would not be expected to contribute to early embryos. To distinguish between these possibilities, we assessed the ability of *Mta123Δ* ES cells to differentiate in chimaeric embryos. Equal numbers of control or *Mta123Δ* ES cells expressing a fluorescent marker were aggregated with wild type morulae and allowed to develop for 48 hours. While wild type cells contributed to the ICM of host embryos with 100% efficiency, *Mta123Δ* ES cells showed significantly reduced contribution and increased levels of apoptosis (Fig. 5B-D). Those *Mta123Δ* cells that did survive in blastocysts were predominantly, but not always found in the inner cell mass and expressed Sox2 but not Cdx2, indicating that they had not undergone inappropriate differentiation towards a trophectoderm fate. This is most consistent with *Mta123Δ* cells being unable to properly adjust to an ICM environment and enter a normal differentiation path. The ability to contribute to the ICMs of chimaeric embryos was rescued by constitutive expression of Mta2 in *Mta123Δ* cells (Fig. 5C). We therefore propose that NuRD functions not only to establish the correct lineage identity of cells during the differentiation process, but also prevents inappropriate gene expression to maintain an appropriate differentiation trajectory (Fig. 6).

**Figure 6.**
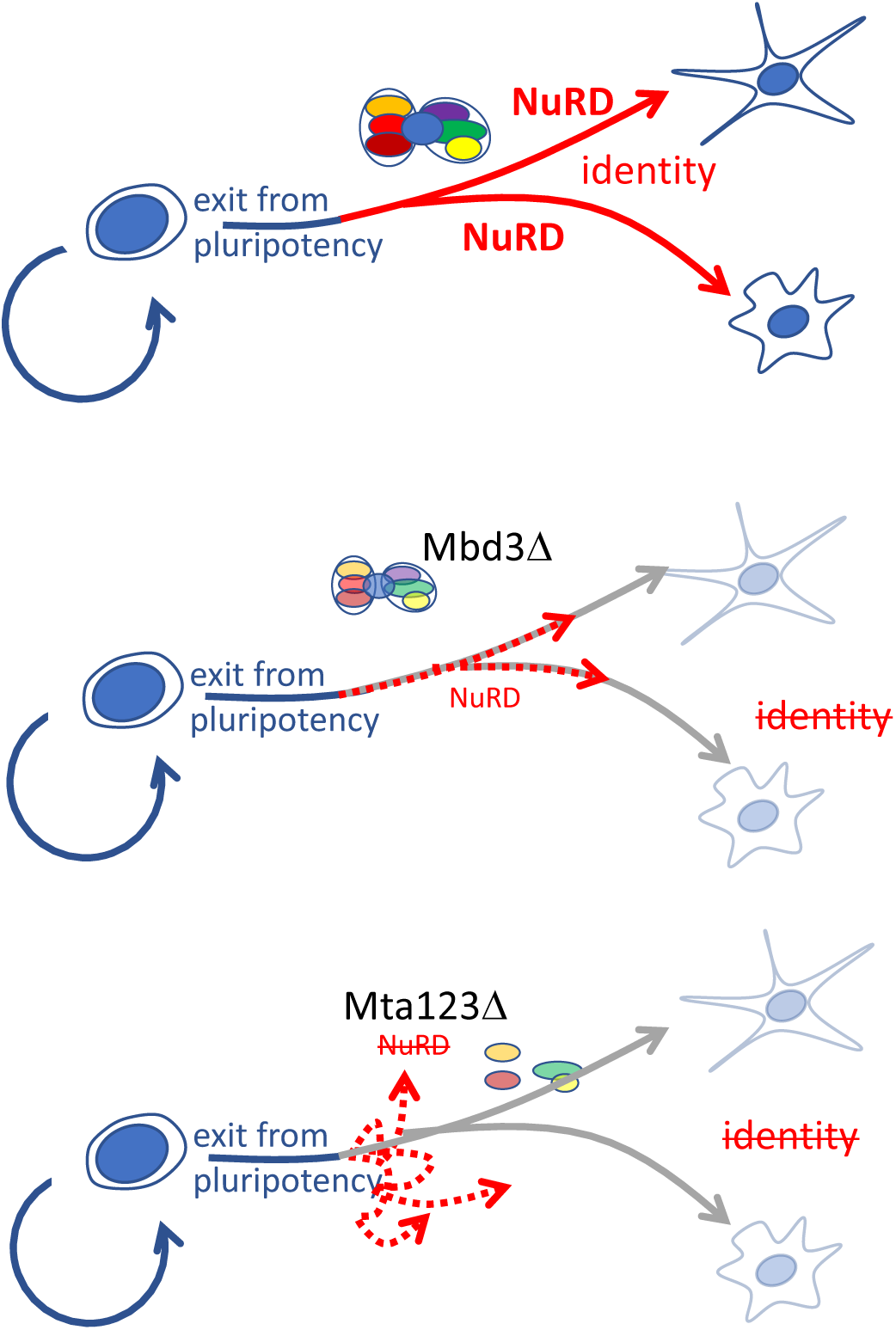
Model of NuRD function during differentiation of pluripotent cells. NuRD facilitates lineage commitment of ES cells after exit from pluripotency (red arrows), allowing cells to form differentiated cell types (Top). In the absence of Mbd3, residual NuRD activity ensures cells retain the appropriate differentiation trajectory, but the cells are unable to reach a differentiated cell fate (dotted red arrows; Middle). In the absence of all three MTA proteins there is no residual NuRD activity and ES cells are unable to either achieve appropriate lineage commitment, or to maintain the proper differentiation trajectory (dotted arrows, bottom panel).

## Discussion

Here we provide a biochemical and genetic dissection of the core NuRD component MTA proteins in mouse ES cells. In contrast to what has been found in somatic cell types, MTA proteins are not mutually exclusive in ES cell NuRD complexes and all combinations of Mta homo- and hetero-dimers can exist within NuRD. Different MTA proteins exhibit subtle differences in chromatin localisation or biochemical interaction partners, but we find no evidence for protein-specific functions in self-renewing or differentiating mouse ES cells. ES cells completely devoid of MTA proteins are viable but show inappropriate expression of differentiation-associated genes, are unable to maintain an appropriate differentiation trajectory and do not contribute to embryogenesis in chimaeric embryos.

Protein subunit diversity is often found in chromatin remodelling complexes which specifies functional diversity (Hargreaves and Crabtree 2011; Morey et al. 2012; Hota and Bruneau 2016). This is also the case for the NuRD complex, where alternate usage of Mbd2/3, Chd3/4/5, and Mta1/3 has been found to result in alternate functions for NuRD complexes (Feng and Zhang 2001; Fujita et al. 2003; Nitarska et al. 2016). Yet our findings that Mta1 and Mta3-null mice are viable and fertile, and that we detect no major differences in the abilities of different MTA proteins to rescue the *Mta123Δ* ES cell phenotypes indicates that MTA proteins exhibit considerable functional redundancy. Different MTA proteins are capable of interacting with each other in ES cells, so how the mutual exclusivity reported in other cell types might be achieved is not clear. One possibility is that the variable inclusion in NuRD of zinc finger proteins, such as the Sall proteins in ES cells, could influence the MTA makeup of NuRD complexes. This class of variable NuRD interactors, which include Sall1/2/3/4, Zfp423 (Ebfaz), Zfpm1/2 (Fog1/2), and Bcl11b, interact with RBBP and/or MTA proteins via a short N-terminal motif (Hong et al. 2005; Lauberth and Rauchman 2006; Lejon et al. 2011). Of this class of proteins Sall1 and Sall4 are the most highly expressed in ES cells, and Sall4 can associate with all three MTA proteins (Fig. 1C) (Miller et al. 2016). In contrast the Zfpm1 (Fog1) protein was shown to preferentially associate with Mta1 and Mta2, but not Mta3, in a somatic cell line (Hong et al. 2005). Hence it is possible that different proteins using this N-terminal motif to interact with NuRD in different cell types could act to skew the proportion of different MTA proteins included in the NuRD complex.

Mta1 and Mta2 both contain two distinct RBBP interaction domains, while Mta3 lacks the C-terminal most RBBP interaction domain (Millard et al. 2016). The Mta1 and Mta2 proteins show minimal loss of stability in the absence of Mbd3, but Mta3 requires an interaction with Mbd3 to be completely stable (Bornelöv et al. 2018). It is possible that the additional interaction with an Rbbp protein confers stability to Mta1 and Mta2 in the absence of Mbd3. This would be consistent with our interpretation that Mta3 preferentially exists within an intact NuRD complex (Figs 1, 2). Rbbp4/7 confer histone H3 binding to the NuRD complex, so Mta3-containing NuRD may bind chromatin less tightly than Mta1- and/or Mta2-containing NuRD. Consistent with this possibility, we identified >2x more Mta1- and Mta2-assoicated ChIP-seq peaks than Mta3-peaks (Fig. 2A).

Pluripotent cells lacking Mbd3 have been used extensively to show that NuRD plays important roles in control of gene expression during early stages of exit from pluripotency in vitro and in vivo (Kaji et al. 2006; Kaji et al. 2007; Latos et al. 2012; Reynolds et al. 2012). Mbd3-null ES cells contain Mbd2/NuRD and thus represent a NuRD hypomorph, rather than a NuRD-null. ES cells lacking all three MTA proteins show no detectable NuRD formation (Fig. 3C) and therefore we propose represent a true NuRD null. *Mta123Δ* ES cells are similar to *Mbd3Δ* ES cells in that both misexpress a range of genes in 2iL conditions and both fail to properly undergo neuroectodermal differentiation. Yet the *Mta123Δ* ES cells misexpress a larger number of genes than do *Mbd3Δ* cells in both self-renewing and differentiation conditions, and they show a more pronounced differentiation defect. We propose that the increased transcription of a wide array of lineage inappropriate genes in *Mta123Δ* ES cells renders them incapable of not only achieving the correct differentiation state, but also of maintaining the proper differentiation trajectory (Fig. 6). In this model the primary function of NuRD in ES cells is to silence inappropriate gene expression, which ensures fidelity of differentiation. A further, more specialised function contained within the core function and dependent upon Mbd3 (but not Mbd2), is to respond properly to context-appropriate differentiation signals to achieve specific differentiation states.

## Materials and Methods

### Mouse embryonic stem cells

Mouse embryonic stem cells (ESCs) were grown on gelatin-coated plates in 2i/LIF conditions as described (Ying et al. 2008). The following “Knockout First” alleles were obtained from EUCOMM as heterozygous ES cell lines (Illustrated in Fig. S1A, B):

#### Mta1^tm1a(EUCOMM)Wtsi^

https://www.mousephenotype.org/data/alleles/MGI:2150037/tm1a(EUCOMM)Wtsi

#### Mta2^tm1a(EUCOMM)Wtsi^

https://www.mousephenotype.org/data/alleles/MGI:1346340/tm1a(EUCOMM)Wtsi

ES cells were used to derive mouse lines by blastocyst injection using standard methods. ESC derivation was performed by isolating ICMs and outgrowing in 2i/LIF ESC media as described (Nichols et al. 2009).

The epitope tagged ESC lines Mta1-Avi-3xFLAG and Mta2-GFP were generated by traditional gene targeting, while the Mta3-Avi-3xFLAG line was generated using a CRISPR/Cas9 genome editing approach. All Mta epitope tagged lines were made in an Mbd3^Flox/-^ background (Kaji et al. 2006). Transient expression of Dre recombinase was then used to remove the selectable marker (Anastassiadis et al. 2009).

ES cells were induced to differentiate towards a neuroectoderm fate by removal of two inhibitors and LIF and culturing in N2B27 media as described (Ying et al. 2008). Mesendoderm differentiation was performed as follows: ES cells were plated at 10^4^ cells/cm^2^ in N2B27 on fibronectin-treated 6-well plates and cultured for 48 hours. Medium was then replaced with 10 ng/ml activin A and 3 μM CHIR99021 in N2B27 and cultured further.

### Mice

All animal experiments were approved by the Animal Welfare and Ethical Review Body of the University of Cambridge and carried out under appropriate UK Home Office licenses. The Mta3 “Knockout First” allele was obtained from EUCOMM as a heterozygous mouse line (Illustrated in Fig. S1C):

#### Mta3^tm3a(KOMP)Wtsi^

https://www.mousephenotype.org/data/alleles/MGI:2151172/tm3a%2528KOMP%2529Wtsi?

Heterozygous knockout first mouse lines were crossed to a Flp-deletor strain kindly provided by Andrew Smith (University of Edinburgh) (Wallace et al. 2007) to generate conditional alleles. Mice harbouring conditional alleles were crossed to a Sox2-Cre transgenic line (Hayashi et al. 2002) to create null alleles.

Chimaeric embryos were made by morula aggregation with 8-10 ES cells per embryo as described (Hogan et al. 1994) and cultured for 24-48 hours prior to fixation and immunostaining. ES cells used stably expressed a PiggyBac Kusabira Orange transgene.

#### Immunoprecipitation

Antibodies were incubated with Protein G-Sepharose beads (Sigma) for 1h at room temperature. Nuclear extract (200 μg) was diluted in IP-Buffer (50 mM Tris-HCl pH 7.5, 150 mM NaCl, 1 mM EDTA, 1% Triton X-100, 10% glycerol) with protease inhibitors and incubated with antibody-bead conjugates at 4 °C overnight. Beads were washed in Low Salt Wash-Buffer (IP-Buffer containing 150 mM NaCl), followed by High Salt Wash-Buffer (IP-Buffer containing 300 mM NaCl). Antibodies are listed in Table S1. Full images of original western blots are available on Mendeley Data: doi:10.17632/sxg8d5sgv6.1.

#### Label-free pulldowns and label-free quantitation (LFQ) LC-MS/MS analysis

Label-free pulldowns were performed in triplicate as previously described (Kloet et al. 2016; Miller et al. 2016). The mass spectrometry proteomics data have been deposited to the ProteomeXchange Consortium via the PRIDE partner repository with the dataset identifier PXD009855.

#### Chromatin immunoprecipitation (ChIP), sequencing, and analysis

Chromatin immunoprecipitations were carried out as previously outlined (Reynolds et al. 2012). For sequencing of ChIP DNA, samples from six (Mta1-FLAG and Mta2-GFP) and four (Mta3-FLAG) individual ChIP experiments were used. Antibodies used are listed in Table S1. ChIP-seq libraries were prepared using the NEXTflex Rapid DNA-seq kit (Illumina) and sequenced at the CRUK Cambridge Institute Genomics Core facility (Cambridge, UK) on the Illumina platform. Reads were aligned using bowtie version 0.12.8 and the arguments −m 1. Peaks were called using macs2 version 2.1.0 with the default Q-value of < 0.05. The results were merged to get 7 different sets of peaks defined by the combination of bound Mta proteins.

DNA Methylation data was obtained from (Shirane et al. 2016). Output methylation files were filtered for base pairs with a coverage of over 3. The methylation percentage was averaged using a sliding window of 250 bp to give a smoothed track for the heatmaps/profile.

High throughput sequence datasets used in this manuscript are listed in Supplemental Table 2.

#### Gene expression analyses

Total RNA was purified using RNeasy Mini Kit (Qiagen) including on-column DNase treatment. First-strand cDNA was synthesized using SuperScript IV reverse transcriptase (Invitrogen) and random hexamers. Quantitative PCR (qPCR) reactions were performed using TaqMan reagents (Applied Biosystems) on a QuantStudio Flex Real-Time PCR System (Applied Biosystems) or a StepOne Real-Time PCR System (Applied Biosystems). Gene expression was determined relative to housekeeping genes using the ΔCt method. TaqMan assays are listed below.

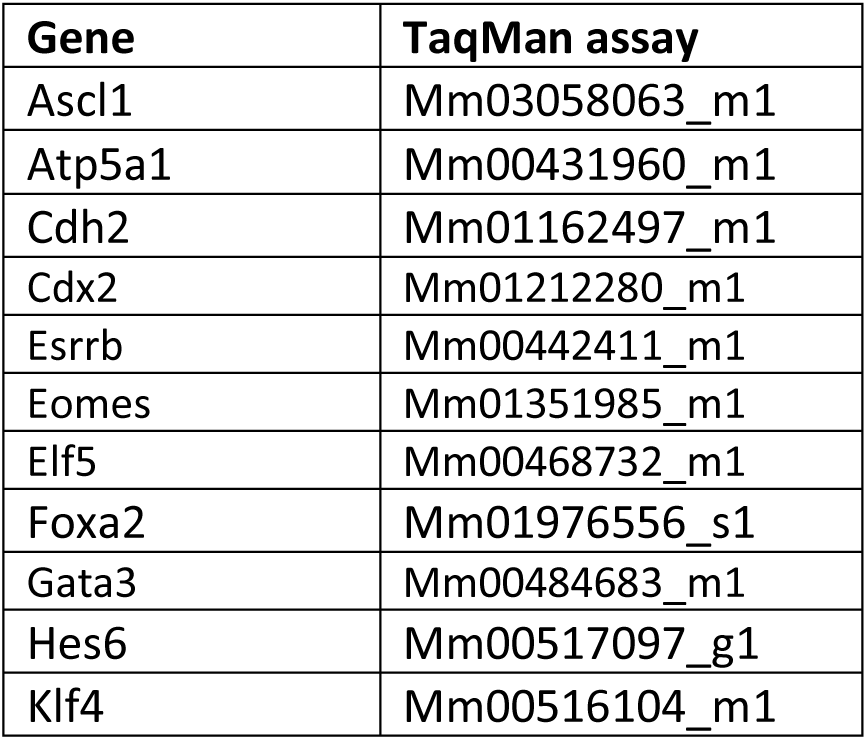

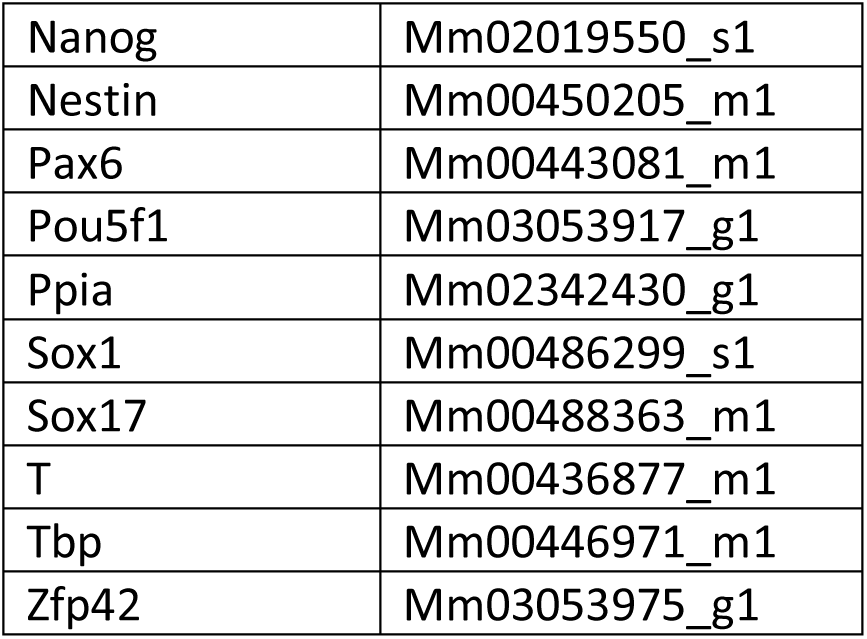

#### RNA-seq

Libraries for sequencing were prepared using the NEXTflex Rapid Directional RNA-seq kit (Illumina) or SMARTer^®^ Stranded Total RNA-Seq Kit v2 - Pico Input Mammalian (Takara Bio) and sequenced on the Illumina platform as for ChIP-seq libraries. Reads were aligned using tophat version v2.1.0 (Kim et al. 2013) to genome build GRCm38/mm10. Gene expression was quantified using featurecount version 1.5.0 (Liao et al. 2014) with annotation from Ensembl release 86 (Yates et al. 2016). Normalization and differential expression were performed using Deseq2 version 1.14.1 (Love et al. 2014), using R version 3.3.3. Deseq size factors were calculated using RNA spike-ins.

## Acknowledgments

We thank Bill Mansfield, Peter Humphreys, Maike Paramor, Vicki Murray, and Sally Lees for technical assistance and advice, and A. Smith, E. Laue and members of the BDH lab for discussions and comments. Funding to the BH and MV labs was provided through EU FP7 Integrated Project “4DCellFate” (277899). The BH lab further benefitted from a Wellcome Trust Senior Fellowship (098021/Z/11/Z) and from core funding to the Cambridge Stem Cell Institute from the Wellcome Trust and Medical Research Council (097922/Z/11/Z and 203151/Z/16/Z).

TB and BH devised the study; TB, SK, SG, RF, NR and BH generated the data; MB, MR and SD analysed high throughput sequencing data, SK and MV generated and analysed proteomics data, MK provided methodology and NR and BH wrote the manuscript with input from other authors.

## Supplemental Data

**Figure S1.**
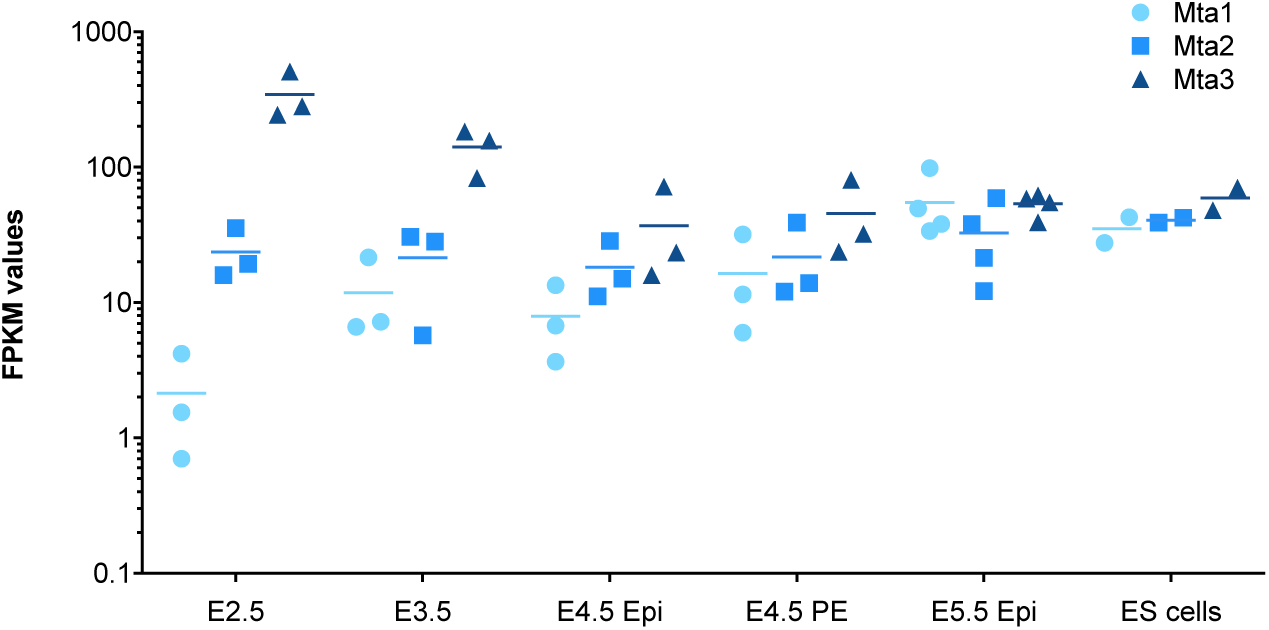
Expression of MTA genes during early mouse development. RNAseq data from (Boroviak et al. 2014) plotted at indicated days of mouse development for each of the MTA genes. All data points are shown, with horizontal bars indicating the mean.

**Figure S2.**
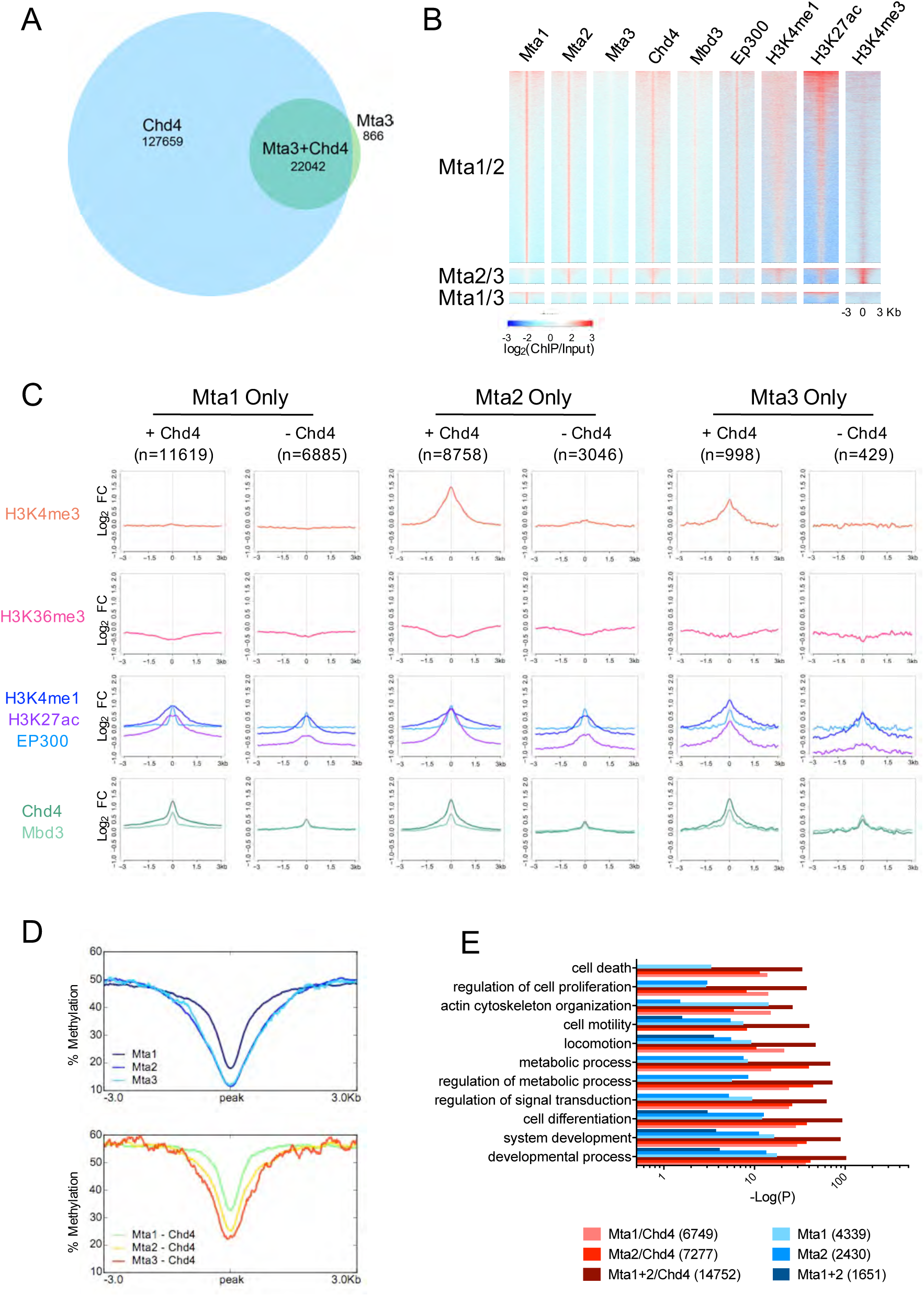
Chromatin features of MTA-bound peaks (related to Figure 2). A. Comparison of ChIP-seq peaks for Mta3 and Chd4. The number of Mta3-only, Chd4-only, or Mta3+Chd4 peaks are indicated. B. Features of peaks containing two of the three MTA proteins, as in Figure 3C. C. Average enrichment of indicated features is shown for peaks for each Mta protein which do or do not also contain Chd4. The number of peaks in each set is indicated as n. D. Average enrichment of DNA methylation across MTA peaks with or without Chd4, as in Panel C. E. Significance of GO-term enrichment for genes associated with peaks for different combinations of Mta proteins and Chd4.

**Figure S3.**
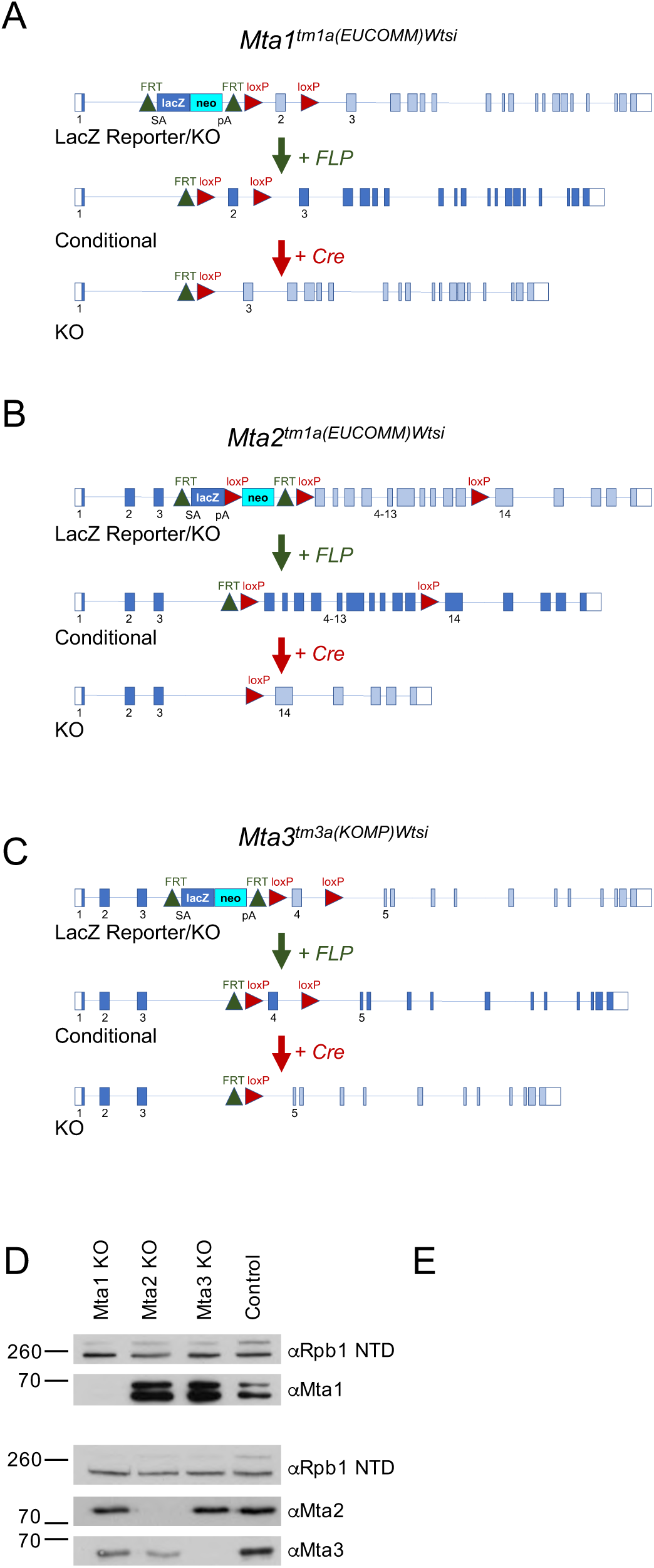
MTA reporter and knockout alleles (related to Figure 3) A. Schematic of the *Mta1* “Knockout First” reporter allele *Mta1^tm1a^*^(^*^EUCOMM^*^)^*^wtsi^* (Top). Exons are depicted as boxes, with normal coding exons as filled boxes. Exons around the insertion site are numbered. Coding exons not able to be translated in the depicted allele are shaded in light blue. The targeting resulted in an FRT-flanked LacZ-Neo fusion protein being expressed from the endogenous *Mta1* promoter and preventing transcription of most *Mta1* exons. Recombination between FRT sites is achieved by expression of FLP recombinase (middle), removing the LacZ-Neo cassette and restoration of Mta1 coding potential. Subsequent recombination between LoxP sites by Cre recombinase (bottom) results in loss of Exon 2 and subsequent exons are out of frame. B. Schematic of the *Mta2^tm1a^*^(^*^EUCOMM^*^)^*^Wtsi^* allele as in panel A. In this allele the neo gene is expressed from a human ß-actin promoter. In this allele expression of Cre results in excision of exons 4-13. C. Schematic of the *Mta3^tm3a^*^(^*^KOMP^*^)^*^Wtsi^* allele as in panel A. D. Western blot of nuclear extracts made from each single mutant (i.e. after Cre-mediated recombination) probed with indicated antibodies. Anti-Rbp1 NTD (RNA polymerase II subunit) acts as a loading control. E. Phase-contrast images of cell lines of indicated genotype in self-renewing conditions. Scale bars indicate 100 μm.

**Figure S4.**
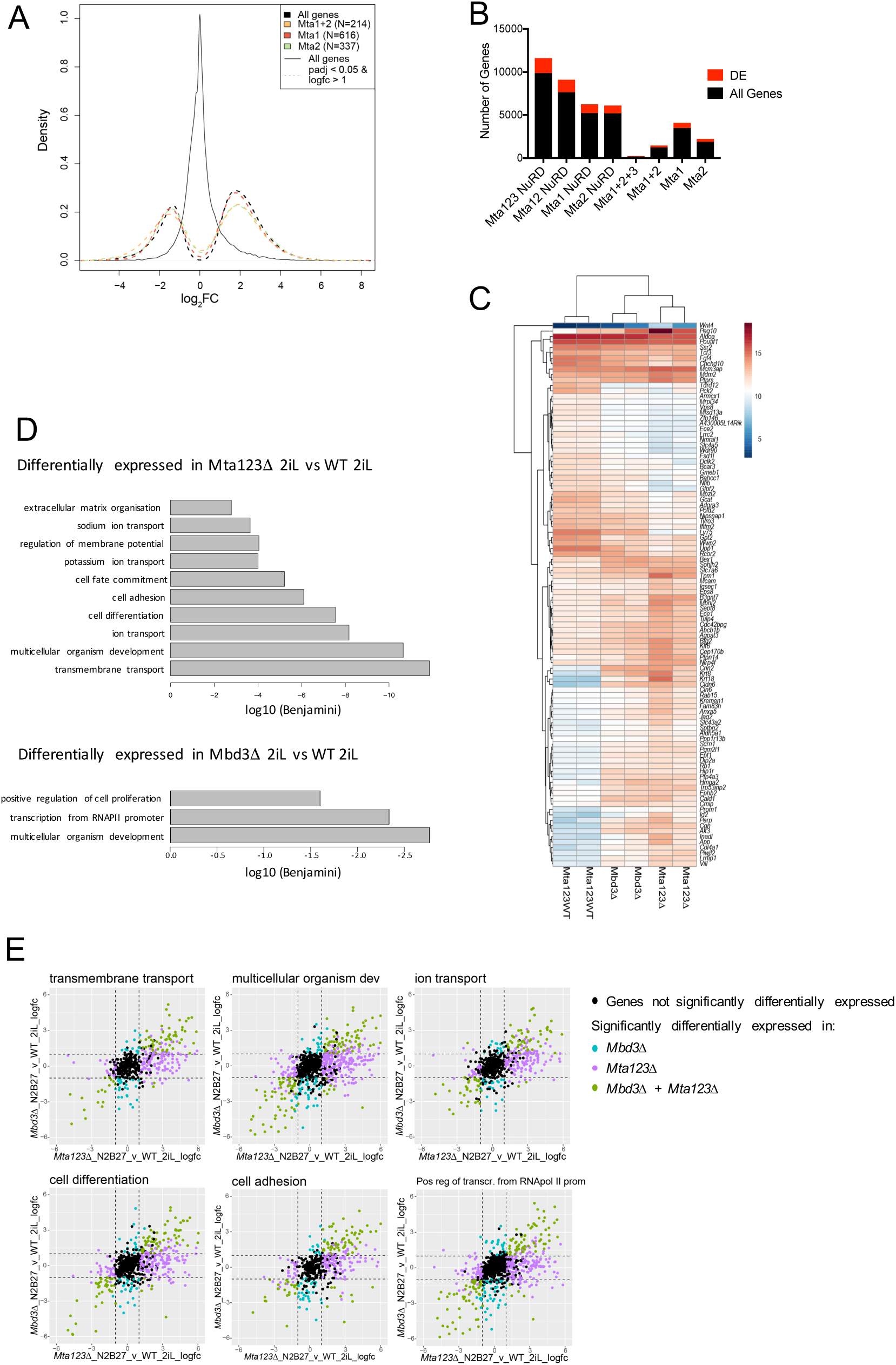
Control of gene expression by the MTA proteins (Related to Figure 4). A. Same as Fig. 5B, but plotting genes associated with Mta peaks lacking Chd4 enrichment. B. Plot showing the number of genes associated with indicated classes of Mta ChIP-peaks which do (red) or do not (black) show a significant change in expression in *Mta123Δ* ES cells in 2iL. C. Comparison of gene expression changes in wild type, *Mbd3-null* (*Mbd3Δ*) and *Mta123* triple-null (*Mta123Δ*) ES cells in self-renewing (2i) conditions. The top 100 genes contributing to PC2 from Figure 5C are shown. “Mta123 WT” indicates a control cell line derived at the same time as the *Mta123Δ* cells. D. GO term enrichment for genes differentially expressed in the indicated comparisons. Significant gene ontology terms are plotted by log_10_ of the Benjamini-adjusted p-value. The significant GO terms and p-values were calculated using David v.6.8 E. Genes associated with indicated GO terms plotted by fold change in expression in *Mta123Δ* ES cells (x-axis) or *Mbd3Δ* ES cells (y-axis) as compared to wild type cells. Each point is a gene that has been annotated with that GO-term. Genes are coloured if they are differentially expressed in either comparison or both, using a log_2_ fold-change > 1 and a padj value < 0.05. The dotted lines are at the fold-change cut-off of 2.

**Figure S5.**
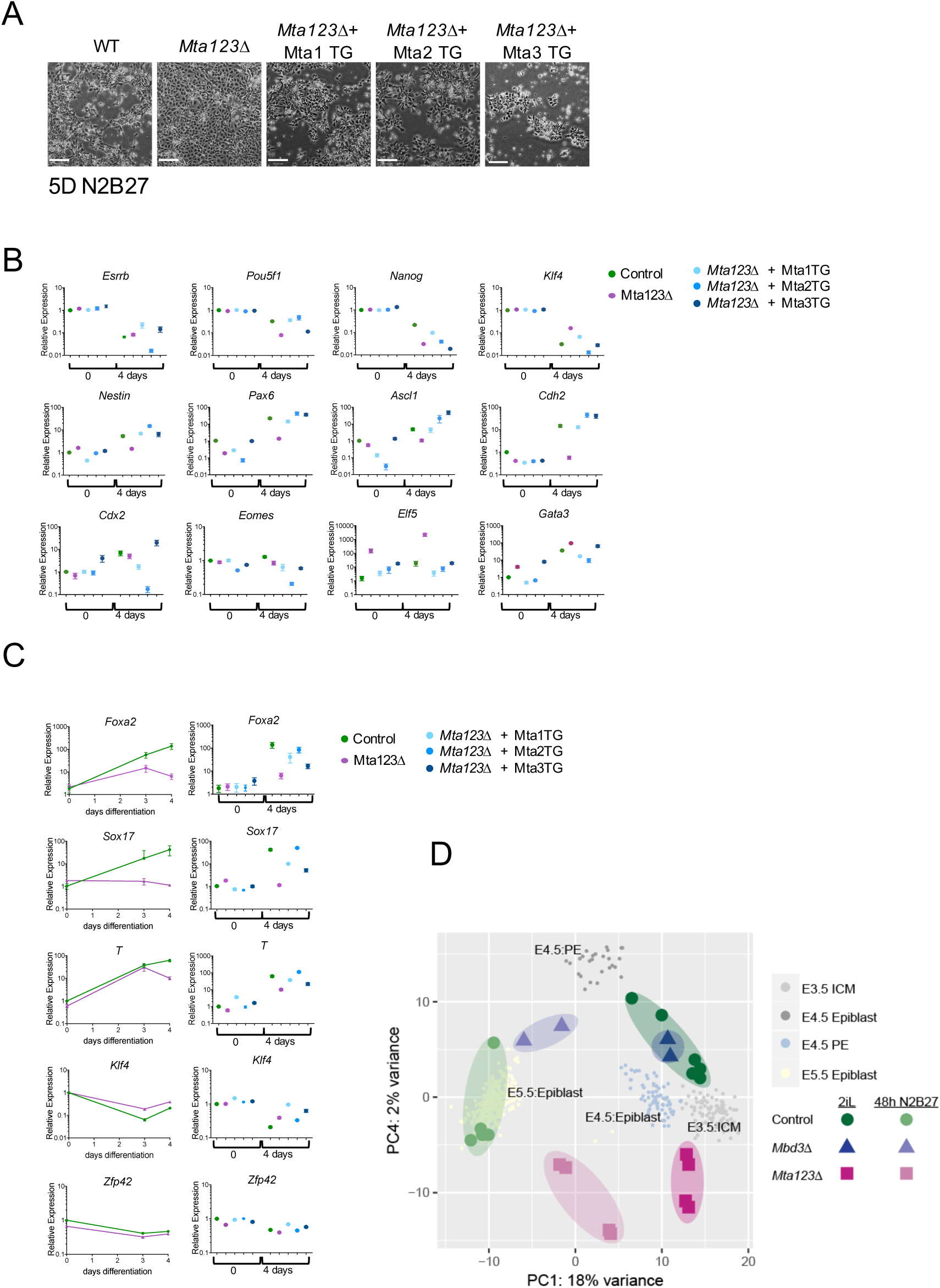
Failure of differentiation in Mta123Δ ES cells (related to Figures 5 and 6). A. Phase contrast images of ES cells of indicated genotypes induced to differentiate for 5 days in N2B27. B. Comparison of gene expression in indicated ES cell lines in undifferentiated conditions or after 4 days differentiation in N2B27. qPCR was carried out in triplicate at each time point for a minimum of three biological replicates. Error bars indicate SEM. C. Comparison of gene expression in indicated ES cell lines induced to differentiate towards mesoderm. qPCR was carried out in triplicate at each time point for a minimum of three biological replicates. Error bars indicate SEM. D. Same as Figure 6A, plotting PC4 vs PC1.

**Table S1.** Antibodies used in this study.

| Antibody | Company<br>(Product Number) | WB | IP | IF | ChIP |
| --- | --- | --- | --- | --- | --- |
| anti-MTA1 | Cell Signaling<br>(5647) | 1:2000 | - |  | - |
| anti-MTA2 | Abcam<br>(ab50209) | 1:5000 | 3 µg |  | - |
| anti-MTA3 | Proteintech<br>(14682-1-AP) | 1:2000 | - |  | - |
| anti-MBD3 | Abcam<br>(ab157464) | 1:5000 | - |  | - |
| anti-CHD4 | Abcam<br>(ab70469) | 1:5000 | 5 µg |  | - |
| anti-HDAC2 | Santa Cruz<br>(sc-7899) | 1:1000 | - |  | - |
| anti-RBBP4 | Abcam<br>(ab79416) | 1:5000 | - |  | - |
| anti-GATAD2B | Bethyl Laboratories<br>(A301-281A) | 1:1000 | - |  | - |
| anti-Histone H3 | Abcam<br>(ab1791) | 1:5000 | - |  | - |
| anti-RNA Pol II<br>subunit RPB1 | Santa Cruz<br>(sc-899X) | 1:2000 | - |  | - |
| anti-FLAG M2 | Sigma-Aldrich<br>(F1804) | 1:2000 | 3 µg |  | 10µg |
| anti-GFP | Abcam<br>(ab290) | 1:5000 | 15µg |  | 25µg |
| anti-Sox2 | e-biosciences<br>(14-9811-82) |  |  | 1:500 |  |
| anti-CASPASE-3<br>cleaved | Cell Signalling<br>(9664S) |  |  | 1:500 |  |
| anti-Cdx2 | Abcam<br>(ab157524) |  |  | 1:250 |  |

**Table S2.** High-throughput sequencing datasets used in this study.

